# Genome-wide analysis of the dynamic and biophysical properties of chromatin and nuclear proteins in living cells with Hi-D

**DOI:** 10.1101/2022.11.17.516893

**Authors:** Cesar Augusto Valades-Cruz, Roman Barth, Marwan Abdellah, Haitham A. Shaban

## Abstract

To understand the dynamic nature of the genome in real-time, the localization and rearrangement of DNA and DNA-binding proteins must be analyzed across the entire nucleus of single living cells. Recently, we developed a new computational light microscopy technique, called high-resolution diffusion mapping (Hi-D), that can accurately detect, classify, and map the types of diffusion and biophysical parameters such as the diffusion constant, anomalous exponent, drift velocity, and physical diffusion models at a high spatial resolution over the entire genome in living cells. Hi-D combines dense optical flow to detect and track local chromatin and protein motion, and Bayesian inference to characterize this local movement at nanoscale resolution. The initial implementation requires solid experience using MATLAB (MathWorks) and computational resources, for instance, access to a computer cluster, to perform the Hi-D analysis. In addition, this implementation takes ∼18-24 hours to analyze a typical imaging stack. To avoid these limitations and emphasize high-performance implementation, we present a customized version called Hi-D-Py. The new implementation is written in the open-source Python programming language and has an option for parallelizing the calculations to run on multi-core CPUs. The functionality of Hi-D-Py is exposed to the users via user-friendly documented Python notebooks. Our efficient implementation reduces the analysis time to less than one hour using a multi-core CPU with a single compute node. We also present different applications of Hi-D for live-imaging of DNA, H2B, and RNA Pol II sequences acquired with spinning disk confocal and super-resolution structured illumination microscopy.

## 1 Introduction

The dynamic organization of chromatin structures is highly relevant to cell function, which may be reflected in the relative distribution of open euchromatin and dense heterochromatin regions^1,2^. According to structural models, the genome is folded into multiple hierarchical levels of chromatin structure, from domain folding and long-range looping to compartments and subcompartments^3^. It is becoming clear that the dynamic nature of these nuclear compartments is involved in the regulation of their organization and thus affects the regulation of nuclear processes^4^. For example, the nucleosome density must be rearranged at transitions within and between euchromatin and heterochromatin to facilitate DNA processing^5^ and coordinated for chromatin-proteins interaction^6,7^. The dynamic activity of chromatin and nuclear proteins in living cells has been studied at high spatiotemporal resolution using specifically labeled molecules that can be identified and tracked in real-time using various techniques^8,9^. Labeled loci and single genes can be tracked over various lengths and time scales using single particle tracking methods^10,11^. However, to fully understand the physical properties of a long-fiber structure, genomic chromatin must be comprehensively studied across the entire nucleus. Nucleus-wide methods achieved quantification of the momentary displacement but these methods neither quantify the extracted motion spatially in terms of physical diffusion models^12,13^ nor at sub-pixel resolution^13^. Therefore, we recently developed the high-resolution diffusion (Hi-D) method^14^. The Hi-D method combines a dense Optical Flow algorithm to quantify the local motion of bulk structures with sub-pixel resolution. Bayesian inference is used to precisely classify types of motion throughout and to assign biophysical parameters, in particular the diffusion constant, anomalous exponent, and drift velocity, to the motion modes throughout the nucleus. Using Hi-D, we have investigated the dynamic roles of chromatin (both DNA and histones)^14–17^, RNA polymerase II^14,18^ heterochromatin protein 1^14^, and transcription factors^19^ in genome organization. Here we present a framework written in Python as an accessible and user-friendly tool, with an improvement in processing time. Using this framework, we present a step-by-step Hi-D analysis of DNA and nuclear proteins imaged with spinning disks confocal and super-resolution microscopy. Documentation and code are released as part of the Hi-D-Py project (https://github.com/haitham-shaban/hidpy).

### 1.1 Applications of Hi-D

Hi-D allows the monitoring of spatiotemporal dynamics within nuclei of living cells in action. Recently, we have shown the applicability of the Hi-D method in studying the dynamic properties and precisely mapping the diffusion parameters and physical diffusion models of chromatin, RNA Pol II, and transcription factors, proteins at nanoscale resolution in single living cells^14,18,19^. We believe that the Hi-D method sets a base for dynamic analysis of chromatin and nuclear macromolecules across many imaging scenarios^14–16^ involving dense signals where well-established methods may reach their limits. Hi-D can potentially be applied to any cellular biomolecule which is sufficiently abundant to dense stain the cell region of interest. Furthermore, we envision that even deep-tissue imaging will be suitable for Hi-D analysis, given a sufficiently high signal-to-noise ratio (>= 20 dB, see SI in^14^). Hi-D has, moreover, shown the capability to be combined with conventional^12,14^ and super-resolution light microscopy techniques^15,16^, and its amenability to screening different cell types. Results from our analysis method open new perspectives toward a global understanding of chromatin organization involved in loop formation^7,20^ potentially the kinetics of gene expression, and a deeper understanding of dynamic processes in many biological fields^8,21^.

### 1.2 Key features and advantages

1. Hi-D is highly effective in analyzing the dynamics of abundant molecules and dense structures directly in single living cells without sacrificing active fluorophore density.
2. There is no need for prior knowledge of sophisticated (sparse) labeling preparations or experience in using advanced microscopy techniques.
3. Commonly used methods to extract diffusion characteristics of nuclear loci such as single particle tracking (SPT) methods cannot be used with dense labeling or certain nuclear regions analysis and require special hardware such as fluorescence correlation spectroscopy (FCS)^22,23^. Hi-D can overcome these limitations because neither sparse labelling nor high imaging are required.
4. Hi-D is adequate to analyze images acquired with both conventional and super-resolution microscopy techniques.
5. Hi-D method uniquely extracts meaningful diffusion characteristics at single pixel resolution since the extracted diffusion is directly related to a physical description of various types of diffusion, by doing so Hi-D overcomes the limitations of averaging nucleus-wide analysis method^13,24^.
6. Hi-D can be applied separately to all kinds of different fluorophores, as long as, those can be spectrally demixed, even in the same cell. This property enables the processing of multi-channel images to spatiotemporally correlate different nuclear factors, as shown in Figure S13 in the original Hi-D article^14^.

### 1.3 Limitations

Hi-D is not a single particle tracking method, so Hi-D does not apply to biological scenarios in which only sparse labeling can be achieved, e.g., due to low protein abundance or where the signal-to-noise ratio is below 20 dB (Figures S7-S8 in the original Hi-D article^14^). Furthermore, small, moving, and particularly bright fluorescence spots might obscure Hi-D analysis as their contribution to the surrounding fluorescence intensity overwhelms the flow field quantification. Hi-D analysis should thus be performed on regions of images without labeling artifacts such as clusters of fluorophores. More extended regions of varying fluorescence intensity do not impose limitations on Hi-D (Figure S7c in^14^). As for all methods quantifying intracellular motion, cells should be de-drifted before Hi-D analysis.

There are technical limitations of the original protocol of the Hi-D method can be summarized in the following points:

1. The protocol was implemented mainly in MATLAB, a commercial software application that requires specific and hefty licensing. The GMM analysis module is implemented using the pomegranate package in Python. The interaction between MATLAB and Python required the user to establish the connection between MATLAB and Python manually, which may limit inexperienced researchers to run the protocol in their research environments.
2. The execution of the current protocol is time-consuming; processing a single video sequence might take around 24 hours or more on mid-range commodity hardware. This issue limits the analysis of lengthy time lapses with specific experimental conditions on personal computers and might require high-end computing clusters for large-scale processing.
3. The existing protocol does not have a flexible interface; it is merely a group of scripts that run independently. The user is required to visualize and compare the results of different cell lines / biological treatments manually. The code was designed as a proof-of-concept to demonstrate the principal idea of the Hi-D method.

### 1.4 Adaptations

We present this protocol to overcome the technical limitations of the existing one by:

1. Re-designing and implementing the workflow in Python, a high-level and non-commercial programming language that does not need any commercial licensing, contrary to MATLAB.
2. Reducing the execution time of the entire workflow by making use of parallel execution environments supported by Python including the multiprocessing and joblib packages by default. Even with a single-threaded execution, the Python implementation is faster than the MATLAB-based one.
3. Making the software more user-friendly and easier to run via documented Python notebooks which guide the user in a step-by-step fashion to analyze the results on the fly.

### 1.5 Overview of the procedure

The full protocol of Hi-D is divided into two complementary procedures. The first procedure is performed in the laboratory to acquire video sequences of living cells showing the dynamics of chromatin and nuclear proteins. Further details on this procedure are available^14^. The second procedure (Stages 1 and 2) analyzes these sequences relying on Hi-D-Py. The presented protocol is focused on the Python workflow implemented for analyzing the video sequences. Nevertheless, we briefly describe a few details of the first procedure for users aiming to replicate the conditions and setups that were used to generate the presented sequences (refer to the video sequences provided within the source code repository and used as default input to the in the jupyter notebooks). The Python workflow is composed of three stages. In Stage 1, the input sequence is pre-processed to compute the Optical Flow fields, estimate the trajectories, and apply Bayesian inference. We then select the most suitable models for deconvolution in Stage 2. In Stage 3, we compare between average statistics of subpopulations from different biological conditions.

#### 1.5.1 Stage 1: Pre-processing, optical flow calculation, trajectories estimation, MSD calculations, and Bayesian inference

After the generation of the video sequence, we applied Stage 1 to pre-process the frames of the input sequence, compute the trajectories and their mean square displacements (MSDs) and use Bayesian inference to test the different models for any MSD curve (Stage 1 in **Figure 1**). The pre-processing stage thresholding, denoising, and edge detection operations are applied to match the specific requirements of the optical flow algorithm. For a sequence with N frames, N-1 Optical Flow maps are produced. Those maps are then interpolated and linked to compute trajectories on a per-pixel basis. Therefore, we allow the user to define a thresholding value based on image intensity to determine if the value of the pixel is significant enough to compute a trajectory or not. The resulting trajectories are then listed in a linear list and mapped to a three-dimensional array for visualization. After trajectory estimation, the MSD curves are calculated for every trajectory. The estimated MSD is a function of the initial position and the time lag (**Supplementary Section 1**). A nucleus mask is estimated during the pre-processing stage and used to select only nuclear localized trajectories. Local MSD estimation is described in the original Hi-D publication^14^.

**Figure 1.**
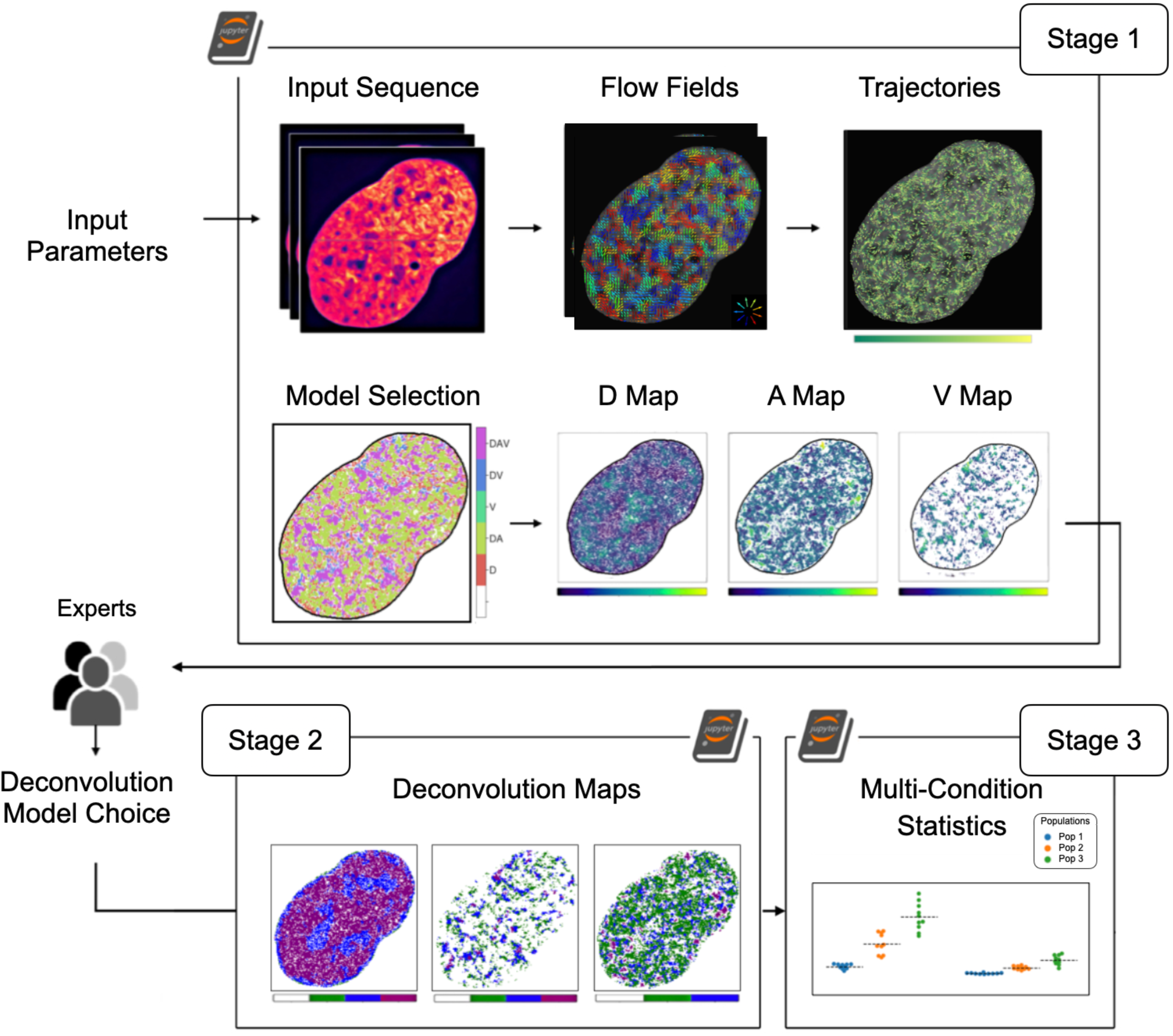
Overview of the Hi-D-Py workflow. This workflow is composed of three stages. In Stage 1, we pre-process the input sequence, compute the flow field and trajectories and then apply Bayesian inference to compute the parameters needed for the deconvolution. Users can read and select the best-fitting models for deconvolution in Stage 2. Stage 3 compares the statistics of subpopulations from different biological conditions.

Bayesian inference is used to test different models for any given MSD curve^25^. Given the data *Y = {Y_1_, …, Y_n_}*, *K* model candidates *M = {M_1_, …, M_K_}* and parameter set *θ = {θ_1_, …, θ_K_}*, the goal is to find the model *M_k_(Y, θ_k_)* such that the probability of *M_k_(Y, θ_k_)* is maximal for the selected set of models; therefore, the optimal parameters for every model are calculated. A general multivariate Gaussian function^26^ is used to represent the probability that the data *Y* is observed for a Model *M_k_* described by the function *M_k_(x; θ_k_)* for any parameter set *θ_k_* as shown in **Supplementary Section 2**. After estimating the best parameter set for a model, the model, and its chosen parameters are selected to maximize their probability to describe the data in which *θ̂*_*k*,*MLE*_= *argP*(*Y*|*θ*_*k*_, *M*_*k*_).

MSD analysis is performed locally^14^, only the 3×3 neighborhood of a pixel is chosen to determine the MSD covariance matrix used for the Bayesian inference^27^. The choice of a window size of 3×3 is based on the selected filter size in the Optical Flow estimation. All the calculations are implemented in Python. Scipy^28^ is used for numerical fitting.

The type of diffusion characterizing each trajectory motion was chosen in an unbiased manner using the Bayesian inference described previously from a set of five common models to fit each trajectory’s MSD. These methods were detailed earlier^14^, and a summary of them is presented in **Table 1**. A constant offset *o* is added to consider experimental noise.

The user can select a subset, or all the models listed in **Table 1**, where D is in units of μm^2^/s. D_α_ is in units of μm^2^/s^α^, α is its anomalous exponent, v [μm/s] is its velocity, and R_C_ [μm] is the radius of a sphere within the particle that is confined. Free diffusion is expected for particle motion in an unrestrictive environment, directed motion may result from an active process or is expected for repelling particles at short time scales. Anomalous diffusion may arise from a variety of factors including diffusion in an environment with obstacles^29^, transient binding events^30^, crowding^31^, or due to the polymeric nature of chromatin^32^. Confined diffusion applies specifically to the analysis of trapped particles, such as membrane proteins^33^. A free, anomalous, and confined motion may co-occur with directed motion.

**Table 1:** Overview of MSD models.

| Model | Equation |
| --- | --- |
| Free Diffusion ($D$) | $\text{MSD}_D(\tau) = 4D\tau + o$ |
| Drift ($V$) | $\text{MSD}_V(\tau) = v^2\tau^2 + o$ |
| Free Diffusion + Drift ($DV$) | $\text{MSD}_{DV}(\tau) = 4D\tau + v^2\tau^2 + o$ |
| Anomalous Diffusion ($DA$) | $\text{MSD}_{DA}(\tau) = 4D_\alpha\tau^\alpha + o$ |
| Anomalous Diffusion + Drift ($DAV$) | $\text{MSD}_{DAV}(\tau) = 4D_\alpha\tau^\alpha + v^2\tau^2 + o$ |
| Confinement + Diffusion (DR) | $\text{MSD}_{DR}(\tau) = R_C^2 \left( 1 - e^{\left( \frac{-4D\tau}{R_C^2} \right)} \right) + o$ |
| Confinement + Diffusion + Drift (DRV) | $\text{MSD}_{DRV}(\tau) = R_C^2 \left( 1 - e^{\left( \frac{-4D\tau}{R_C^2} \right)} \right) + v^2\tau^2 + o$ |

#### 1.5.2 Stage 2: Deconvolution by a General Mixture Model

The Bayesian inference of the trajectory MSD results in maps of the diffusion constant (*D*), anomalous exponent (*A*), and drift velocity (*V)* for every cell (the best model parameters *θ̂*_*k*,*MLE*_). These parameter values are represented in histograms and subsequently deconvolved by a General Mixture Model (GMM) approach^27^. Before starting the trajectory computation and Bayesian inference (Stage 2 in **Figure 1**), the user selects the maximum number of sub-populations *n_ot_* to be expected within the data set, as well as the type of distribution to be used by the GMM. Without prior knowledge, the number of sub-populations can be set to 3, and typically both normal and log-normal distributions can be considered. The GMM is then evaluated on all types of distributions for a number of sub-populations from 1 to *n_pop_*. The goodness of fit for each of those combinations is evaluated using the Bayesian information criterion^34^ (*BIC*) (Supplementary Section 3). The result of the deconvolution is a table representing which fractions of cells in the analyzed data set are best described by a particular combination of distribution type and the number of sub-populations for every parameter (*D*, *A*, and *V*). The user should inspect this table (as shown in **Figure 2**). In principle, every cell might be best described by a different number of sub-populations of varying relative size and different shapes (i.e., normal, or log-normal). To enable a global GMM deconvolution across the entire data set, the user is advised to select the GMM settings from the table which describes the largest fraction of cells within the dataset; this should be the default choice. Currently, Hid-py considers this automatically. However, if the user has reason to believe that another setting would be more appropriate to describe the data, potentially due to prior knowledge, the user may select another setting. The table is then used for the user to evaluate which fraction of cells within the dataset is well described by a particular choice of GMM settings. When a decision has been reached, the user could leave default choice or to supply the number of subpopulations and shapes and starts the final deconvolution stage with the chosen model(s) to obtain deconvolved histograms as well as a mapping of the respective distributions within the cell nucleus. If the Jupyter notebook is used, the parameters are updated in the parameters panel directly.

**Figure 2.**
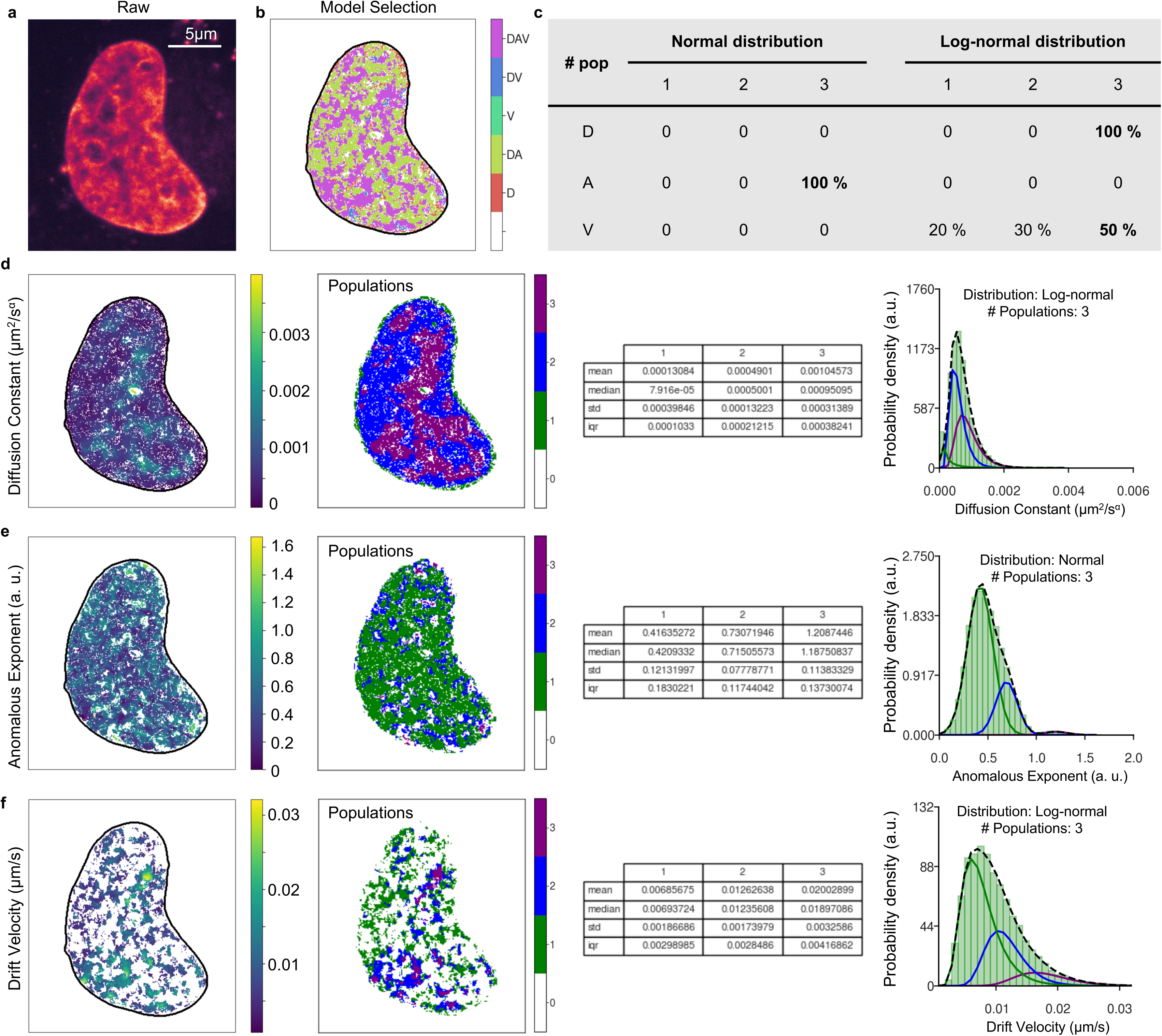
Deconvolution by a General Mixture Model (GMM) approach in Hi-D-Py. SiR-DNA-stained U2OS cells with Serum were imaged at Δt = 200 ms. Optical Flow estimation and Bayesian inference were done using Hi-D-Py. Representative maps are shown for one cell: raw (**a**) and model selection (**b**). (**c**) A table displaying the distribution type and the number of sub-populations for each parameter (diffusion constant (D), anomalous exponent (A), and drift velocity (V)) within the dataset (n=10 cells). The optimal combination is highlighted in bold numbers for each parameter. (left **d-f**) Representative coefficient maps are shown for one cell. An estimated populations’ map (center-left **d-f**) and histograms (right **d-f**) were estimated using the optimal combination for parameter D. Coefficient statistics for each population are shown in (center-right **d-f**).

#### 1.5.3 Stage 3: Comparison between average statistics of subpopulations from different biological conditions

In this stage the users can compare between the biophysical and diffusion parameters out of the Hi-D-py analysis from different biological conditions (Stage 3 in **Figure 1**). The three stages are described in detail in Section 3, in order to assist the user to use the protocol in step-by step procedure.

### 1.6 Evaluation of the protocol implementation in Python

Dense Optical Flow fields are reconstructed by an Optical Flow algorithm based on the Horn-Schunck formulation^35^. The performance of the translation to python was evaluated against the previously used MATLAB implementation^14,36^ (Figure S1) on experimental data from the original work. The MATLAB implementation partially utilizes built-in functions for which no source code is available. We carefully validated that the python translation returns identical results to the MATLAB implementation for all available source code. Nevertheless, the translation of proprietary algorithms yields slightly different results, which ultimately results in small variations of the estimated flow fields (endpoint error median ±. SD = 0.67 ± 0.39 pixels, ratio between flow magnitude of the python and MATLAB implementation median ± SD = 0.996 ± 1.967; **Figure S1**). **Figure S2** shows a comparison between the results of the Bayesian analysis^1^ using MATLAB and that resulting from our Python implementation. Similar results are shown for the least-square fitting from MATLAB Curve fitting toolbox and Python Scipy library.

### 1.7 Installation · TIMING a few minutes

The code is installed in a few easy steps as follows:

1. Go to the open-source GitHub repository at https://github.com/haitham-shaban/hidpy.
2. If you have a GitHub account, clone the repository to download its contents into the file system. **CRITICAL STEP** If you wish to contribute to the code, you can fork the repository before cloning it.
3. If you do not have a GitHub account, you can download the project as a . Zip file.

### 1.8 Code execution · TIMING 1 hour to 5 hours (depends on the CPU cores available in your machine)

Hi-D-Py can be easily executed from a list of Python notebooks, allowing the user to run every stage interactively. All the notebooks are documented and labeled, making the connections between all the stages seamless. The code consists of three principal notebooks 01-hidpy-stage-1.ipynb, 02-hidpy-stage-2.ipynb, and 03-hidpy-stage-3.ipynb. The 00-install-dependencies.ipynb contains a single panel that allows the user to verify if the installed Python environment where the notebooks are executed has all the needed dependencies to successfully run all the code. Note that the code supports parallel execution using all available CPU cores in your computing node as executed from the jupyter notebooks. We also added detailed documentation of how to use the code in a step-by-step fashion on the Wiki page of the online repository.

## 2 Materials

### 2.1 Biological materials

A list of all the biological materials including the cell lines, chemicals, and relevant data is summarized in Table 2.

**Table 2:** Key resource table for all the biological materials used in this protocol.

| Resource or reagent | Source | Identifier |
| --- | --- | --- |
| Dulbecco's Modified Eagle Medium, high glucose, pyruvate | ThermoFisher | Cat#41966029 |
| Fetal Bovine Serum | ThermoFisher | Cat#10270106 |
| Sodium pyruvate solution | Sigma-Aldrich | Cat#113-24-6 |
| Penicillin-Streptomycin | Bioconcept | Cat#4-1F00H |
| Puromycin | Thermofisher | Cat#A1113803 |
| Glutamax | ThermoFisher | Cat#35050061 |
| Trypsin-EDTA-Solution | Sigma-Aldrich | Cat#T4049 |
| Doxycycline hyclate | Sigma-Aldrich | Cat#D9891 |
| SiR-DNA (5-610CP-Hoechst) | Grazvydas Lukinavicius | N/A |
| L-15 Medium (Leibovitz) | ThermoFisher | Cat#11415064 |
| HEPES buffer | Sigma-Aldrich | Cat#7365-45-9 |
| Live-cell images of DNA, H2B, and RNA pol II data (Spinning disk confocal microscopy). | Shaban et al., Genome Biology, 2020 | Reference <sup>14</sup> |
| Live-cell images of H2B data (Structured illumination microscopy). | Miron et al., Science Advances, 2020 | Image Data Resource (IDR0086) |
| U2OS cell line | ATCC | N/A |
| U2OS cell line stably expressing H2B-GFP | ATCC. A gift from Sébastien Huet (Rennes) | N/A |

### 2.2 Equipment

1. Spinning disk confocal microscopy system (CSU-X1-M1N, Yokogawa). This system is used to image live-DNA 150-frames sequences with an exposure time of 200 ms per frame. Other imaging modalities like widefield microscopy can be used, but users should be aware that the resulting dynamics might be slightly altered at low signal-to-noise levels below 20 dB (Fig. S9 in^14^).
2. A Nipkow-disk confocal microscopy system (CSU22, Yokogawa). This system is used to image live H2B and RNA Pol II 150-frames sequences with an exposure time of 200 ms per frame.
3. Super-resolution structured illumination microscopy (DeltaVision OMX V3 Blaze). This system is used to image 20 frames of live H2B sequences with a frame duration of 2 seconds. The system is equipped with a PlanApo 60/1.42 oil immersion objective, a 488 nm laser, and sCMOS cameras.
4. Standard personal computer. All protocols have been tested on three machines with different operating systems to ensure that our implementation is cross-platform. The reference machine runs Linux (Ubuntu 20.04) and has an Intel Corei9 CPU with 2.8 GHz and 128 GB of DDR4 memory. Hi-D-Py can be downloaded and run locally as a stand-alone application on Windows, Linux, and macOSX machines.

### 2.3 Software

1. Scientific computing package: OpenCV (opencv.org) and its Python bindings.
2. Python’s scientific computing package: Numpy (numpy.org).
3. Python’s scientific computing package: Scipy (scipy.org).
4. Python’s plotting package: Matplotlib (matplotlib.org).
5. Python’s plotting package: Seaborn (seaborn.pydata.org).
6. Python’s probabilistic models’ package: Pomegranate (pomegranate.readthedocs.io).
7. Python’s pipelining package: joblib (joblib.readthedocs.io).
8. Python’s concurrency package: multiprocessing (docs.python.org/3/library/multiprocessing.html).
9. Jupyter notebook with Visual Studio Code (https://code.visualstudio.com).

All the Python packages can be directly installed by relying on the pip command. The code has been tested on Unix-based operating systems (Ubuntu 20.04 and macOSX 11) and Windows 10. It is recommended to use Visual Studio Code (code.visualstudio.com) to run the Python notebooks.

## 3 Procedure

As a characteristic example, a full Hi-D-Py workflow was applied to a spinning disk confocal sequence of H2B-GFP-stained nuclei of a U2OS cell (see **Figure 3a**). First, trajectories estimated from **Stage 1** are shown in **Figure 3b**. Then, Bayesian inference was applied. Hi-D-Py generates maps of diffusion models (see **Figure 3c**) and the diffusion constant, anomalous exponent, and drift velocity (and the radius of confinement, depending on the set of permitted MSD models; see **Figure 3d**). Finally, deconvolution by a General Mixture Model was applied to the biophysical parameters (*D*, *A*, *V*). The result of this deconvolution stage is a particular combination of distribution type and the number of sub-populations for every parameter. Histograms and maps of populations are shown in **Figures 3e, and f**, respectively.

**Figure 3.**
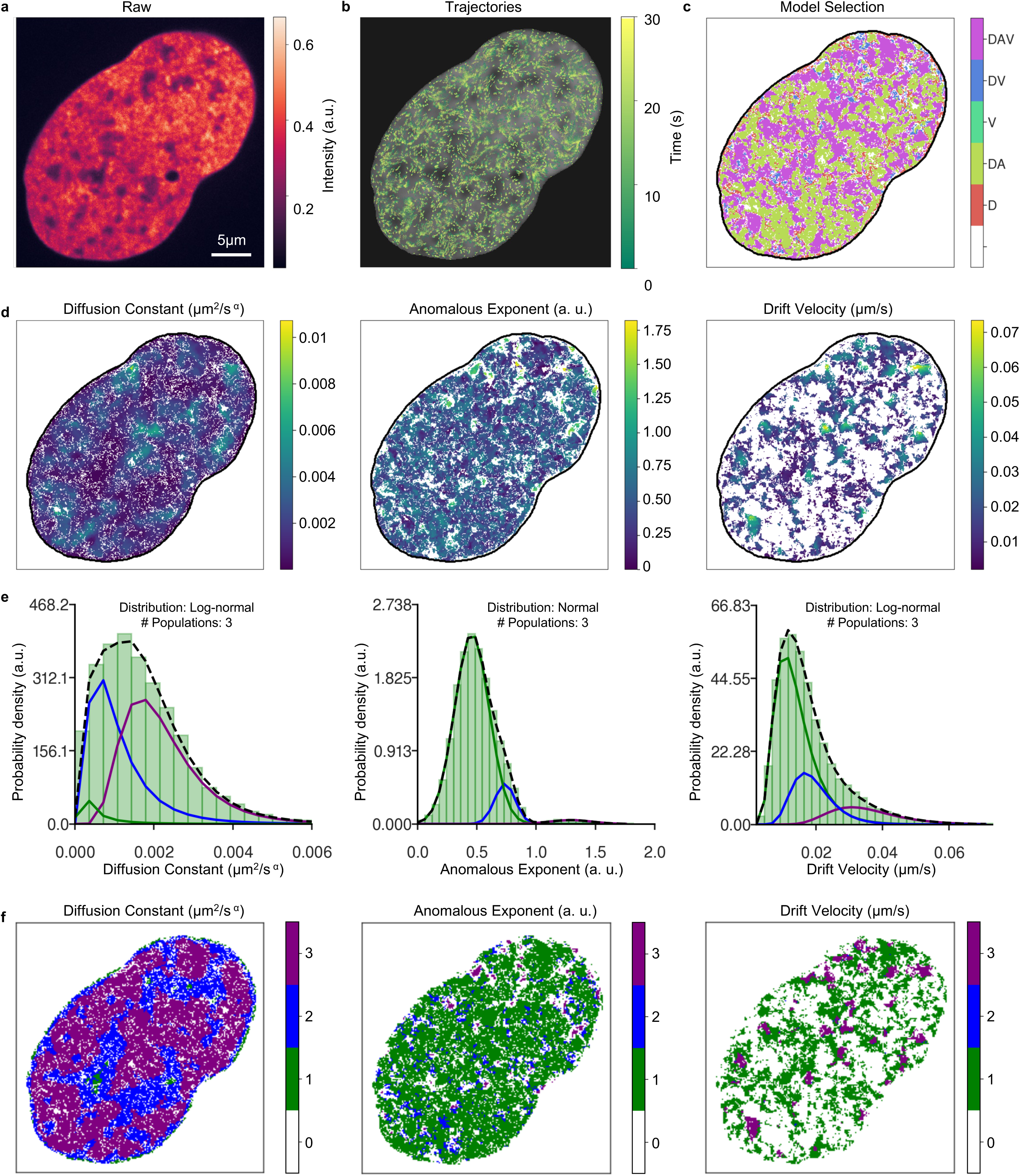
A complete analysis example using the Hi-D-Py. The workflow was applied to a sequence of H2B (**a**) in U2OS with a time interval of Δt = 200 ms. Trajectories were estimated (**b**) and Bayesian inference was used to map the results for the models (**c**) and biophysical parameters (diffusion constant (left **d**), anomalous exponent (center **d**), and drift velocity (right **d**)). Results from deconvolution using GMM are shown, with histograms in (**e**) and the maps of populations in (**f**).

### 3.1 Stage 0: Cell culturing, staining and live-cell imaging, and sequence generation

1. *Cell-culturing and staining.* For H2B imaging using spinning disk confocal microscopy, a human osteosarcoma U2OS cell line stably expressing green fluorescent protein (GFP) tagged histones (H2B) was used. The same cell line U2OS that expresses RPB1 (subunit of RNA Pol II) fused with Dendra2 is used for RNA Pol II imaging, as previously described in^37^. Using an incubator at 37°C with 5% CO2, cells were grown in Dulbecco’s modified Eagle’s medium containing phenol red-free, Glutamax with 50 g/ml gentamicin, 1 mM sodium pyruvate, 10% fetal bovine serum and G418 with 0.5 mg/ml. At a density of about 1×10^5^ cells per petri dish (#1.5 coverslip-like bottom), cells were plated for one day before the imaging. For H2B imaging using super-resolution structured illumination microscopy, HeLa H2B-GFP cells were seeded using phenol red-free DMEM enhanced with 10 M Hepes buffer. For DNA staining of the cultured same cell line, the SiR-DNA (5-610CP-Hoechst) dye was used. On the day of imaging, SiR-DNA was introduced to the medium of the cultured cells at 37°C for an hour after at a final concentration of 1 M. The cells were then gently washed three times with prewarmed phosphate-buffered saline. Before the nuclei imaging, the medium was changed to Leibovitz’s L-15 Medium adding all the medium supplements needed for live cell imaging. This step was performed as well for cells that express H2B-GFP, RNA Pol II-Dendra2, and DNA-SiR for live chromatin imaging.
2. *Live-H2B and RNA Pol II imaging using spinning disk confocal microscopy.* The microscope CSU22 (Yokogawa) was used to record a 150-frame sequence with an exposure time per frame of 200 ms. A diode-pumped solid-state laser with a single wavelength of 488 nm (25 mW) was used to image the H2B-GFP. For RNA Pol II (Rbp1-Dendra2) imaging, a laser source (5-10% light intensity) with a wavelength of 488 nm was applied. Imaging was performed with a Plan Apo (100×/1.42 NA) oil immersion objective from Nikon and detected using a cooled electron-multiplying charge-coupled device camera (iXon Ultra 888), with a sample pixel size of 88 nm. The imaging was performed in a humid chamber at 37°C.
3. *Live-DNA imaging using spinning disk confocal microscopy.* The microscope CSU-X1-M1N (Yokogawa) was used to record a 150-frame sequence with an exposure time per frame of 200 ms of DNA-SiR. For DNA-SiR molecules imaging, an excitation laser source with 140 mW power and a wavelength of 647 nm was used. The emitted light was collected with a Leica HCX-PL-APO (100×/1.4 NA) oil immersion objective lens. A CMOS camera (ORCA-Flash4.0 V2) with (1×1 binning) was used to record the imaged videos with a sample pixel size of 65 nm. The imaging was performed in a humid chamber at 37°C.
4. *Live-H2B imaging using super-resolution structured illumination microscopy (SIM).* A human (HeLa) somatic cell line stably expressing GFP tagged H2B was imaged using SIM. The microscope DeltaVision OMX V3 Blaze was used to record a 20-frame series with an interval between frames of 2 seconds with a sample pixel size of 41 nm. At 37°C and 5% CO2 supply, live-cell imaging was carried out utilizing an objective heater and stage-top incubator.

### 3.2 Stage 1: Pre-processing, Optical Flow calculation, trajectories construction, MSD estimation, and Bayesian inference · TIMING one to five hours per image sequence

1. CAUTION We recommend downloading the provided datasets (**Supplementary Data**) and using the Jupyter notebooks as a *step-by-step* guide to get an understanding of the code with a validated example. A full example is shown in **Figure 3** for analysis of H2B in U2OS, data is available in (**Supplementary Data**).
2. Select and crop single nuclei and convert the input video sequence into a TIFF file format. TROUBLESHOOTING.
3. The input parameters are set in the input panel of the notebook 01_hidypy_stage-1.ipynb. TROUBLESHOOTING.
4. *Denoising*. Each frame of the entire sequence is denoised using non-iterative bilateral filtering to verify the difference in intensity values to preserve the edges of the structures.
5. *Optical flow computations.* Compute the optical flow fields of the entire sequence. For a video sequence of N frames, a list of N-1 Optical Flow fields is computed.
6. *Trajectory estimation.* Starting from the first frame, compute the trajectories of every pixel whose value is above a given intensity threshold defined by the user. Trajectory computations are implemented without taking into consideration the dimensions of the pixel. The pixel size is defined by the user in the input data before running the protocol based on the configuration of the microscope used to image the sequence. Therefore, we scale each estimated trajectory by multiplying the computed coordinates by the pixel size to convert from pixel units to micrometers. We then use the Scipy library to interpolate the trajectories.
7. *Visualize the resulting trajectories.* Map the estimated trajectories on the first frame of the sequence to verify that the trajectories are located within the extent of the nucleus. TROUBLESHOOTING
8. *Trajectory maps reconstruction.* Once the estimated trajectories are calculated and verified, they are converted to flow fields maps (in microns). TROUBLESHOOTING.
9. *MSD estimation.* For each trajectory inside the nucleus mask, the MSD is estimated.
10. *Bayesian inference.* Bayesian inference^14,25^ is applied to trajectories and thereby the best-fitting model is estimated considering the model type and the corresponding parameters (*D*, *A*, *V*). CAUTION This step requires the time step (Δt) between the frames in the input sequence. This step should be set manually by the user depending on the imaging procedure and the configuration of the microscope.
11. *Model selection and parameters are mapped*. The resulting maps of diffusion parameters are the optimal parameters of the MSD model which have been chosen by the Bayesian selection scheme and are thus parameters from a single optimal fit. Examples of these mappings are shown in **Figure 3c,d**. Additionally, a map of coefficient of determination (R^2^) from least-square fitting is included for inspection **(Figure S3**). The default value is R^2^ > 0.6. A higher R^2^ threshold implies higher statistical significance, but it could also lead to a reduction in the number of analyzed pixels (**Figure S3**). It is possible to implement custom diffusion models, we refer the experienced user to the source code file (hidpy/core/inference.py/msd_fitting()). The diffusion constant D’ is also shown as the generalized diffusion constant *D*^′^ = log_10_(*D* ⋅ 1 *s*^*⍺*^/*μm*^2^)^38^, which is a dimensionless quantity, in **Figure S4** TROUBLESHOOTING.
12. *Storing the Bayesian results.* The Bayesian results for all the parameters (*D*, *A*, and *V*) are stored in a pickle file for processing in the next stage.
13. Stage 1 is implemented for a single sequence. Batch execution could be performed using the Python library Papermill, as indicated in the Hi-D-Py wiki (Batch Analysis). The stage 2 and 3 could be run in multiple files as described in the following steps.

### 3.3 Stage 2: Deconvolution by a general mixture model · TIMING a few seconds to a minute

14. *Setting deconvolution parameters.* Before GMM deconvolution is performed, users must choose the maximum number of distributions (*n_pop_*), the model parameter to deconvolve and the type of distribution (normal or log-normal) to be used by the GMM. The input to the GMM step is the raw values resulting from the Bayesian inference stage **(Stage 1**). Model parameter values are represented in histograms before deconvolution. CAUTION The number of histogram bins is selected by the user for visualization purposes. If you are using notebooks, set the input parameters of the pickle files directory and models in the input parameters panel of the notebook 02-hidypy-stage-2.ipynb. TROUBLESHOOTING.
15. *GMM evaluation*. GMM is evaluated on all types of distributions and for subpopulations (1 to *n_pop_*). Evaluation is done using the Bayesian information criterion (*BIC*). The deconvolution results are summarized in a table representing the fraction of cells from the analyzed dataset. The results are best described by a combination of distribution type and the number of sub-populations for every model parameter.
16. *GMM table evaluation.* Users should inspect the GMM table and select the optimal combination of distribution type and the number of sub-populations for each parameter within the analyzed data set. Unless previous knowledge about the number of populations and/or the distribution is available, the user is advised to choose the number of populations and distribution type which encompasses the largest fraction of cells in the dataset.
17. *Deconvolution*. GMM deconvolution with the chosen model(s) is performed and population distributions are mapped within the cell nucleus for each selected model parameter. The final results from GMM deconvolution in the example workflow are shown as histograms (see **Figure 3e**) and maps (see **Figure 3f**) for each biophysical parameter. TROUBLESHOOTING.

### 3.4 Stage 3: Comparison between average statistics of Subpopulations from different biological conditions

18. GMM deconvolution generates a pickle file named Statistics.pickle for each analyzed folder. It is recommended to generate one Statistics.pickle file per condition that will be compared.
19. *Set comparison parameters.* Prior to performing the comparison, users should create a list containing the paths of each Statistics.pickle file for each condition in the input panel of notebook 03-hidypy-stage-3.ipynb. Additionally, the names of each condition and the list of deconvolved parameters should be defined.
20. *Comparison*. Once the input panel is set, execute 03-hidypy-stage-3.ipynb. This will generate a swarmplot that compares the different conditions and populations. An example output from this stage is shown in **Figure 4**. In this evaluation, SiR-DNA-stained U2OS cells, both serum-starved and serum-stimulated (n=10 cells per condition), were used, as described in the original Hi-D publication^14^. Identically as the original Hi-D results^14^, the diffusion constant (V) and anomalous exponent (A) values of DNA in serum-starved U2OS nuclei showed the same changing patterns in all three populations upon the addition of serum (**Figure 4a, b**). Similarly, drift velocity of DNA in serum-starved U2OS nuclei decreased in all three populations upon the addition of serum (**Figure 4c**).

## 4 Troubleshooting

Troubleshooting recommendations for anticipated issues are summarized in **Table 3**.

**Table 3:** Troubleshooting table.

| Step | Problem | Possible Reason | Solution |
| --- | --- | --- | --- |
| 2 | Input video sequences are not loaded. | Unsupported file format. | Convert the sequence into any of the supported sequences, for example, .tiff stack. |
| 3 | Results are not generated to the specified output directory. | The directory is not valid. | Verify the path of the output directory. |
| 4 | Noisy video sequences after bilateral denoising. | Imaging artifacts such as noise. | Use any image processing software, for example: Fiji, to denoise the sequence using median filters. The window size of the bilateral denoising might be increased by 2 pixels. |
| 7 | Trajectories are appearing outside the locus of the nuclei. | Imaging artifacts such as noise. | Re-adjust the pixel threshold value and repeat Steps 1 – 10 until a convenient mapping is reached. The acceptable range of the threshold is between 0 and 1, and the default value is 0.05. The value can only be adjusted by trial-and-error due to several factors including the imaging conditions and the nature of noise in the images, etc. |
| 7 | Trajectories all move collectively in a certain direction. | Nucleus contains drift | Drift may be eliminated using Fiji or another image processing program. Care should be taken to correct lateral drift smoothly and only use integer pixel values to avoid interpolation of pixel's intensity which alters pixel intensity values and often blurs images. |
| 11 | Parameters are higher than expected. | Error in units. | Verify pixel size and time step in input configuration. The units for the pixel size are micrometers. |
| 11 | Bayesian maps show a region outside of the cell. | Error in pixel threshold. | Adjust the pixel intensity threshold value. |
| 11 | Execution failure in case of long or high-resolution videos. | Memory limitations. | Use a single core for processing instead of multiple cores. |
| 11 | Fitting failure in Bayesian inference. | Fitting parameters are not adapted. | Correct fitting parameters in the Python file core/inference.py |
| 14 | Error empty sequence in deconvolution stage. | Empty pickle folder. | Verify that at least one pickle file is in the folder given to the deconvolution stage. |
| 17 | Execution failure during GMM constrained. | The model fitting did not converge. | Try different numbers of populations. and/or distribution type. |

**Figure 4.**
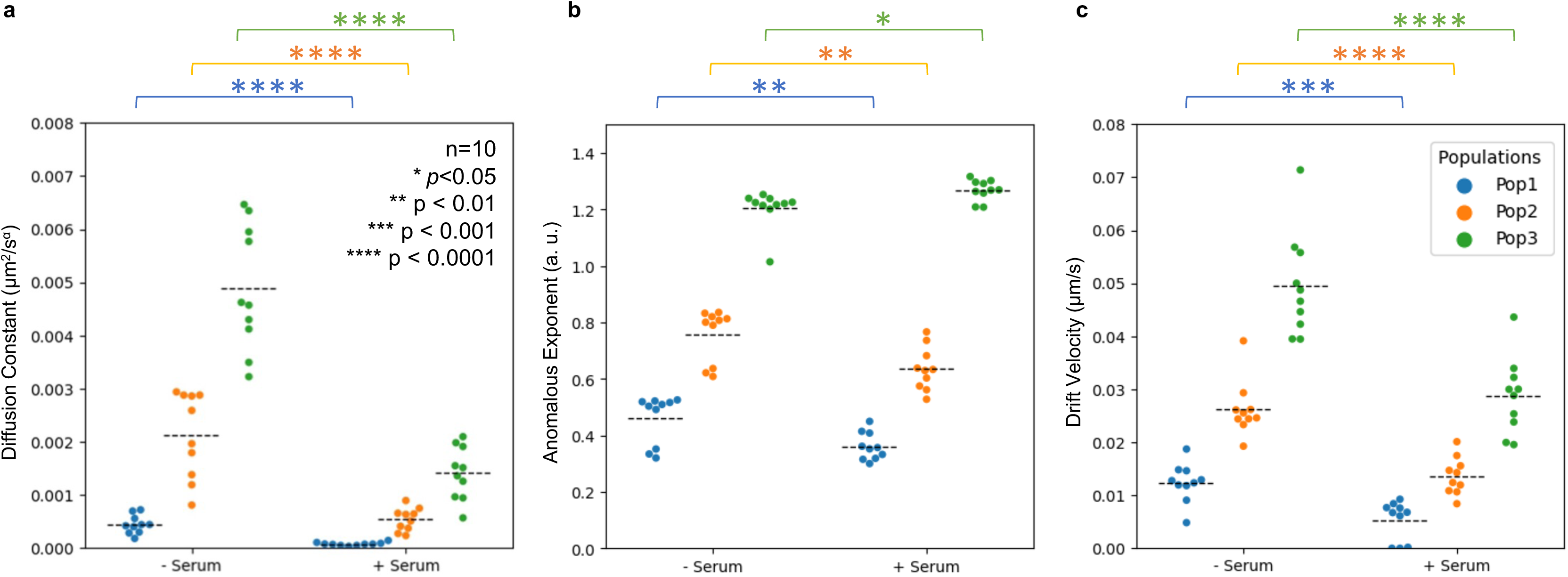
Comparison between two experimental conditions using Hi-D-Py. SiR-DNA-stained U2OS cells, both serum-starved and serum-stimulated (n=10 cells per condition), were analyzed. Comparisons of average values for diffusion constants (**a**), anomalous exponents (**b**) and drift velocities (**c**) per population and per cell are shown. Statistically, a t-test was performed using the Python library scipy.

## 5 Timing

The timing reported in the current protocol is registered based on several video sequences with 150 frames. However, we expect a total running time – on average– between 1 hour to 5 hours to run the entire pipeline. This timing is computed based on the hardware described in the Materials sections. The processing time of the Optical Flow and Bayesian inference steps depends on the number of cores in the CPU if the multicore functionality is enabled. This assumes that the running machine has sufficient memory if lengthy video sequences are supplied. The timing is estimated under the assumption that the user has basic coding experience in Python and understands how to run Python notebooks either via Jupyter Notebooks or relying on Visual Studio Code (Microsoft). Average timing estimates for the entire protocol based on the exemplar datasets are listed in Table 4.

**Table 4:** Average timing estimates for the entire protocol.

| Time estimates for the development and execution of protocol steps |  |
| --- | --- |
| Process | Execution |
| Data Preprocessing | Several seconds to a few minutes. |
| Optical Flow computations | 10-100 s per frame |
| Trajectory computations | A few minutes. |
| MSD-Bayes Analysis | A few to several minutes. |
| GMM Deconvolution | A few seconds. |

The timing also depends on the following factors:

- *Video length.* While the protocol is applied to video sequences with 150 frames, it is assumed that it can run with input sequences with thousands of video frames as long as it has a high signal-to-noise ratio, negligible photobleaching, and corrected lateral drift.
- *Pixels per frame.* The computations of the optical flow fields depend on the number of pixels per frame.
- *Pixel threshold value.* Increasing the value of the pixel threshold will reduce the number of calculated trajectories.
- *Selected models.* Bayesian inference implies a fitting process per trajectory for each selected model, population number, and type of statistic distribution. Execution time is highly dependent on the combination of these factors.

## 6 Anticipated results

The Hi-D-Py software provides an open-source improved implementation of the former Hi-D algorithm. This method permits the quantification, classification, and mapping of diffusion parameters and physical diffusion models over the entire genome in living cells. Hi-D-Py takes advantage of the Python programming environment to improve speed and user-friendliness through Jupyter Notebooks. Hi-D-Py has been applied to two different microscopy techniques showing versatility. First, it has been validated on structured illumination microscopy for the analysis of H2B (see **Figure 5a-d**). Second, it has also been tested on a spinning disk for the analysis of DNA (**Figure 5e-h**) and RNA (**Figure 5i-l**) The different mobility classes of the diffusion constant, anomalous exponent, and drift velocity reflect the cell’s dynamic landscape. Outputs generated by Hi-D-Py may be used to determine the impact of a certain biological condition (a drug interfering with nuclear processes or external stimuli) on a cell to determine mobility differences between cell lines. The found number of populations is informative about e.g., the extent of chromatin-bound, chromatin-processing, and unbound fractions of a nuclear molecule. Similarly, chromatin (re)-arrangements and different complications that associated different cell states and pathogenic conditions can be correlated to chromatin mobility^21^. The mean/median values of *D*, *A*, and *V* per sub-population allow us to express mobility changes quantitatively. We thus encourage the chromatin organization community to complement the presented Hi-D method with external stimuli and additional markers of nuclear processes of interest, both locus-specific, as well as genome-wide. The incorporation of Hi-D-Py as a simple, fast, and user-friendly analysis package for nuclear mobility greatly enhances the aspects of nuclear biology research beyond static and population-average methods as well as restrictive sparse loci methods.

**Figure 5.**
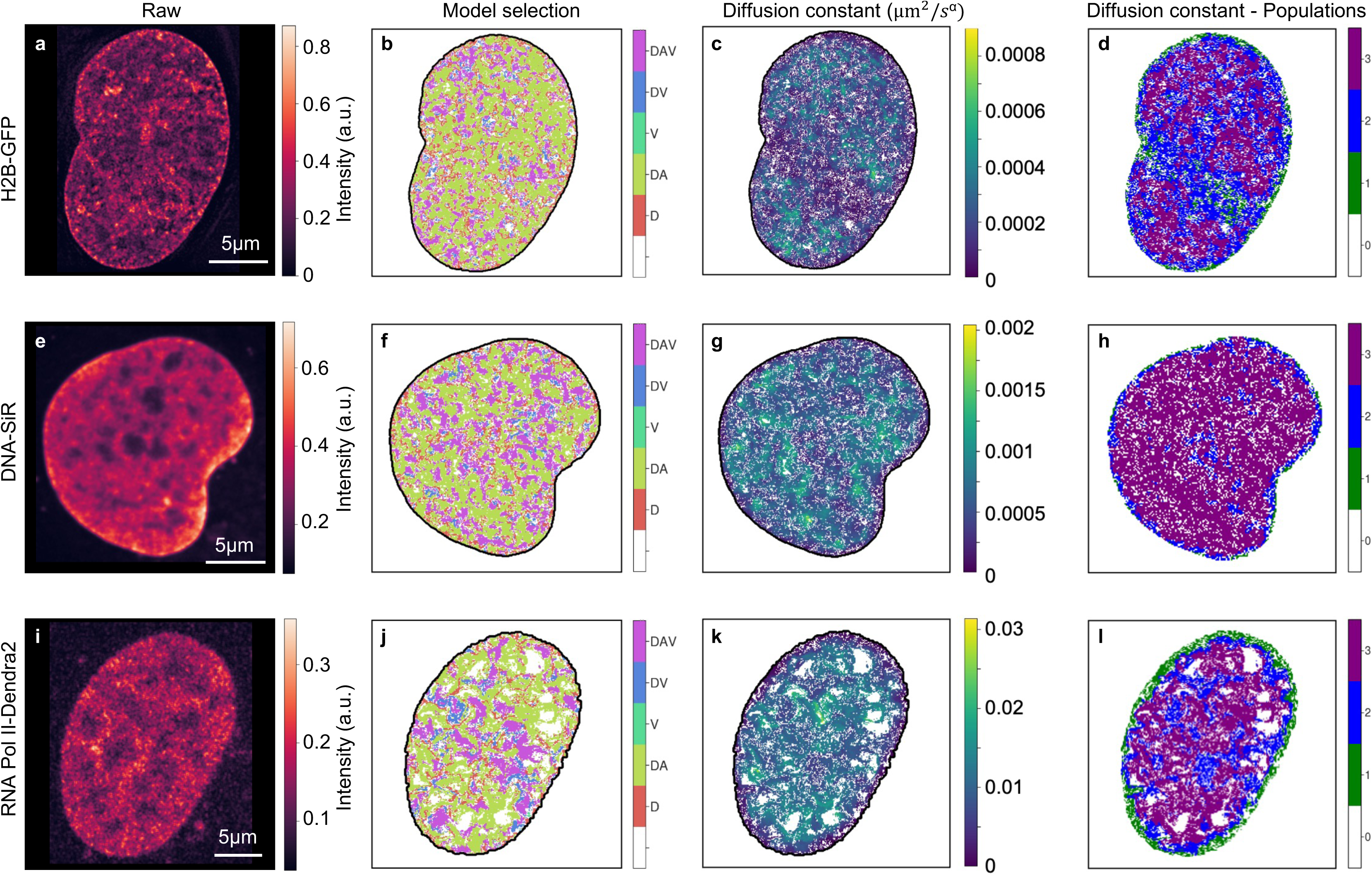
Hi-D-Py applied to analyze live cell imaging of chromatin and nuclear proteins using two microscopy modalities. (**a - d**) Live-nucleus imaging of Hela Cell expressing H2B-GFP imaged using SIM with Δt = 2s. Live cell imaging of DNA-SiR stained (**e - h**) and RNA Pol II (Rbp1-Dendra2) expression in U2Os cells (**i - l**) imaged using a spinning disk microscope with Δt = 0.2s. Representative maps are displayed for each sample type: model selection (**b, f, j**), diffusion constant (**c, g, k**), and the estimated population map (**d, h, l**) from deconvolution using GMM are shown for the analyzed cells H2B, DNA, and RNA Pol II, respectively.

### 6.1 Software availability

The code is open source and freely available online on GitHub at https://github.com/haitham-shaban/hidpy. The version of the code that was used to create the results demonstrated in tagged **v.1** on the GitHub repository. The code in this protocol has been peer-reviewed by all the co-authors.

## Documentation

Documentation is available on GitHub at https://github.com/haitham-shaban/hidpy/wiki.

## Supplementary Data

The example datasets that are used in this protocol are available at https://github.com/haitham-shaban/hidpy.

## Acknowledgments

H.A.S. acknowledges the financial support of the ISREC foundation. C.A.V-C acknowledges the funding by Labex Cell(n)Scale (ANR-11-LABX-0038) as part of the Idex PSL (ANR-10-IDEX-0001-02). M.A. and R.B. contribute to this work independently.

## Authors’ contributions

H.A.S. co-conceived the study. C.A.V-C and M.A. implemented the core framework. R.B. implemented the Optical Flow and validation modules. C.A.V-C. implemented the Bayesian and deconvolution modules. M.A. implemented the trajectory computations. H.A.S. performed the live-cell experiments and imaging protocols. C.A.V-C., R.B., M.A., and H.A.S wrote the manuscript. C.A.V-C. and R.B. contributed equally to this work. H.A.S. supervised the project. All authors approved the manuscript.

## Correspondence and requests for materials

should be addressed to H.A.S.

## References

1 Misteli T. The Self-Organizing Genome: Principles of Genome Architecture and Function. Cell. 2020. doi:10.1016/j.cell.2020.09.014.

2 Agbleke AA, Amitai A, Buenrostro JD, Chakrabarti A, Chu L, Hansen AS et al. Advances in Chromatin and Chromosome Research: Perspectives from Multiple Fields. Mol. Cell. 2020. doi:10.1016/j.molcel.2020.07.003.

3 Dekker J, Mirny L. The 3D Genome as Moderator of Chromosomal Communication. Cell 2016; 164: 1110–1121.

4 Bhat P, Honson D, Guttman M. Nuclear compartmentalization as a mechanism of quantitative control of gene expression. Nat Rev Mol Cell Biol 2021; 22: 653–670.

5 Klemm SL, Shipony Z, Greenleaf WJ. Chromatin accessibility and the regulatory epigenome. Nat. Rev. Genet. 2019. doi:10.1038/s41576-018-0089-8.

6 Shaban HA, Barth R, Bystricky K. Navigating the crowd: visualizing coordination between genome dynamics, structure, and transcription. Genome Biol. 2020. doi:10.1186/s13059-020-02185-y.

7 Gabriele M, Brandão HB, Grosse-Holz S, Jha A, Dailey GM, Cattoglio C et al. Dynamics of CTCF- and cohesin-mediated chromatin looping revealed by live-cell imaging. Science (80-) 2022; 376: 496–501.

8 Shaban HA, Seeber A. Monitoring the spatio-temporal organization and dynamics of the genome. Nucleic Acids Res 2020. doi:10.1093/nar/gkaa135.

9 Shaban HA, Seeber A. Monitoring global chromatin dynamics in response to DNA damage. Mutat. Res. - Fundam. Mol. Mech. Mutagen. 2020. doi:10.1016/j.mrfmmm.2020.111707.

10 Germier T, Kocanova S, Walther N, Bancaud A, Shaban HA, Sellou H et al. Real-Time Imaging of a Single Gene Reveals Transcription-Initiated Local Confinement. Biophys J 2017; 113: 1383–1394.

11 Gu B, Swigut T, Spencley A, Bauer MR, Chung M, Meyer T et al. Transcription-coupled changes in nuclear mobility of mammalian cis-regulatory elements. Science (80-) 2018. doi:10.1126/science.aao3136.

12 Shaban HA, Barth R, Bystricky K. Formation of correlated chromatin domains at nanoscale dynamic resolution during transcription. Nucleic Acids Res 2018; 46: e77.

13 Zidovska A, Weitz D a, Mitchison TJ. Micron-scale coherence in interphase chromatin dynamics. Proc Natl Acad Sci U S A 2013; 110: 15555–60.

14 Shaban HA, Barth R, Recoules L, Bystricky K. Hi-D: nanoscale mapping of nuclear dynamics in single living cells. Genome Biol 2020; 21: 95.

15 Miron E, Oldenkamp R, Brown JM, Pinto DMS, Xu CS, Faria AR et al. Chromatin arranges in chains of mesoscale domains with nanoscale functional topography independent of cohesin. Sci Adv 2020. doi:10.1126/sciadv.aba8811.

16 Barth R, Bystricky K, Shaban HA. Coupling chromatin structure and dynamics by live super-resolution imaging. Sci Adv 2020; 6. doi:10.1126/sciadv.aaz2196.

17 Barth R, Fourel G, Shaban HA. Dynamics as a cause for the nanoscale organization of the genome. Nucleus 2020; 11: 83–98.

18 Barth R, Shaban HA. Spatially coherent diffusion of human RNA Pol II depends on transcriptional state rather than chromatin motion. Nucleus 2022; 13: 194–202.

19 Shaban HA, Suter DM. Individual activator and repressor transcription factors induce global changes in chromatin mobility. bioRxiv 2022. doi:10.1101/2022.04.12.488001.

20 Mach P, Kos PI, Zhan Y, Cramard J, Gaudin S, Tünnermann J et al. Cohesin and CTCF control the dynamics of chromosome folding. Nat Genet 2022; 54: 1907–1918.

21 Shaban HA, Gasser SM. Dynamic 3D genome reorganization during senescence: defining cell states through chromatin. Cell Death Differ 2023. doi:10.1038/s41418-023-01197-y.

22 Hinde E, Cardarelli F, Digman MA, Gratton E. In vivo pair correlation analysis of EGFP intranuclear diffusion reveals DNA-dependent molecular flow. Proc Natl Acad Sci 2010; 107: 16560–16565.

23 Di Bona M, Mancini MA, Mazza D, Vicidomini G, Diaspro A, Lanzanò L. Measuring Mobility in Chromatin by Intensity-Sorted FCS. Biophys J 2019; 116: 987–999.

24 Eshghi I, Eaton JA, Zidovska A. Interphase Chromatin Undergoes a Local Sol-Gel Transition upon Cell Differentiation. Phys Rev Lett 2021; 126: 228101.

25 Monnier N, Guo S-M, Mori M, He J, Lénárt P, Bathe M. Bayesian approach to MSD-based analysis of particle motion in live cells. Biophys J 2012; 103: 616–26.

26 Seber, G. & Wild C. Nonlinear regression. John Wiley & Sons, 2003.

27 Schreiber JM, Noble WS. Finding the optimal Bayesian network given a constraint graph. PeerJ Comput Sci 2017; 2017: 1–16.

28 Virtanen P, Gommers R, Oliphant TE, Haberland M, Reddy T, Cournapeau D et al. SciPy 1.0: fundamental algorithms for scientific computing in Python. Nat Methods 2020; 17: 261–272.

29 Saxton MJ. Anomalous diffusion due to obstacles: a Monte Carlo study. Biophys J 1994; 66: 394–401.

30 Saxton MJ. Diffusion of DNA-Binding Species in the Nucleus: A Transient Anomalous Subdiffusion Model. Biophys J 2020; 118: 2151–2167.

31 Banks DS, Tressler C, Peters RD, Höfling F, Fradin C. Characterizing anomalous diffusion in crowded polymer solutions and gels over five decades in time with variable-lengthscale fluorescence correlation spectroscopy. Soft Matter 2016; 12: 4190–4203.

32 Nieman GC, Robinson GW. Rapid triplet excitation migration in organic crystals. 1962 doi:10.1063/1.1733439.

33 Saxton MJ, Jacobson K. Single-particle tracking: applications to membrane dynamics. Annu Rev Biophys Biomol Struct 1997; 26: 373–399.

34 Wit E, van den Heuvel E, Romeijn JW. ’All models are wrong. ’: An introduction to model uncertainty. Stat Neerl 2012; 66: 217–236.

35 Sun D, Roth S, Black MJ. A quantitative analysis of current practices in optical flow estimation and the principles behind them. Int J Comput Vis 2014; 106: 115–137.

36 Farnebäck G. Two-frame motion estimation based on polynomial expansion. Lect Notes Comput Sci (including Subser Lect Notes Artif Intell Lect Notes Bioinformatics) 2003. doi:10.1007/3-540-45103-x_50.

37 Cisse II, Izeddin I, Causse SZ, Boudarene L, Senecal A, Muresan L et al. Real-Time Dynamics of RNA Polymerase II Clustering in Live Human Cells. Science (80-) 2013; 341: 664–667.

38 Benelli R, Weiss M. Probing local chromatin dynamics by tracking telomeres. Biophys J 2022; 121: 2684–2692.

