## Supplementary Information for "Genome-wide analysis of the dynamic and biophysical properties of chromatin and nuclear proteins in living cells with Hi-D"

#### Supplementary Notes

##### 1 Mean Square Displacement estimation

The mean square displacement (MSD) curves are calculated for every trajectory according to

$$MSD(\overrightarrow{r_0}, \tau) = \langle \left| \overrightarrow{\xi_{r_0}}(t+\tau) - \overrightarrow{\xi_{r_0}}(t) \right|^2 \rangle$$

where $\overrightarrow{\xi_{r_0}}(t)$ represents the position at time $t$ of a particle with initial position $\overrightarrow{r_0}$. $\tau$ are time lags where $\Delta t$ is the time difference between subsequent frames and the average $< \cdot >_t$ is calculated over time. The estimated MSD is a function of the initial position and the time lag.

The analytical solution of the MSD of anomalous diffusion ($DA$) and directed motion ($V$) are calculated as

$$MSD_{DA}(\tau) = 4D_\alpha \tau^\alpha$$
$$MSD_V(\tau) = v^2\tau^2$$

where $D_\alpha$ represents the diffusion constant (in $\mu m^2/s^\alpha$), $\alpha$ is the anomalous exponent (free diffusion: $\alpha = 1$, anomalous diffusion: $0 < \alpha < 1$ and super-diffusion: $1 < \alpha < 2$), $v$ (in $\mu m/s$) is the velocity[1].

##### 2 Bayesian inference of MSD curves

Given the data $Y = \{Y_1, ..., Y_n\}$, $K$ model candidates $M = \{M_1, ..., M_K\}$ and parameter set $\theta = \{\theta_1, ..., \theta_K\}$, the goal is to find the model $M_k(Y, \theta_k)$ such that the probability of $M_k(Y, \theta_k)$ is maximal for the selected set of models, therefore, the optimal parameters for every model are calculated. A general multivariate Gaussian function[2] is used to represent the probability that the data $Y$ is observed for a Model $M_k$ described by the function $M_k(x; \theta_k)$ for any parameter set $\theta_k$:

$$P(Y|\theta_k, M_k) = \frac{1}{\sqrt{(2\pi)^2 \, det(C)}} \cdot exp\left\{-\frac{1}{2}[Y - M_K(x;\theta_K)]^T \cdot C^{-1} \cdot [Y - M_K(x;\theta_K)]\right\}$$

where $C$ is the covariance matrix of the data and the prefactor is a normalizing factor. After estimating the best parameter set for a model, the model and its chosen parameters are selected to maximize their probability to describe the data in which $\hat{\theta}_{k,MLE} = \arg\max_{\theta_K} P(Y|\theta_k, M_k)$.

#### 3 Deconvolution

The goodness of fit for each of those combinations is evaluated using the Bayesian information criterion ($BIC$)[3]:

$$BIC = k \ln(n) - 2 \ln(L)$$

where $k$ denotes the number of parameters per combination of distribution type and number of sub-populations, $n$ denotes the number of data points, and $L$ denotes the maximized value of the likelihood function.

#### 4 Evaluation of the Optical Flow implementation in python.

The endpoint error ($EE$) is computed as

$$EE = |\boldsymbol{V} - \boldsymbol{V}'|$$

where $\mathbf{V}$ denotes the estimated flow field by the python, $\mathbf{V}^{'}$ the estimated flow field by the MATLAB implementation, and $|\cdot|$ denotes the absolute value. We further quantify the flow field magnitude ratio between the python and MATLAB implementations as

$$\text{ratio} = \langle(\boldsymbol{V} - \boldsymbol{V}')/\boldsymbol{V}'\rangle$$

where $\langle\cdot\rangle$ denotes the average over the x- and y-component of the flow fields.

Figure S1 shows a comparison between the original MATLAB code of Optical Flow and python implementation of the Optical Flow module. The translation was validated stepwise on the same input image sequence in both algorithms. The MATLAB implementation made use of proprietary code in two instances (an image interpolation and a Sobel edge detector scheme), whose output could not be exactly reproduced with python implementations. However, we implemented/chose the python implementations that most closely resemble the MATLAB implementation. Even though the numerical difference is small, the Optical Flow algorithm repeatedly uses an image/flow field interpolation step in a pyramidal scheme, which may potentially amplify small numerical differences. We thus compared the final output of both algorithms in Figure S1. The endpoint error has a median value of $0.67 \pm 0.39$ px (median $\pm$ SD) while the ratio has a median value of $0.99 \pm 2.21$ px (median $\pm$ SD).

#### Supplementary Figures

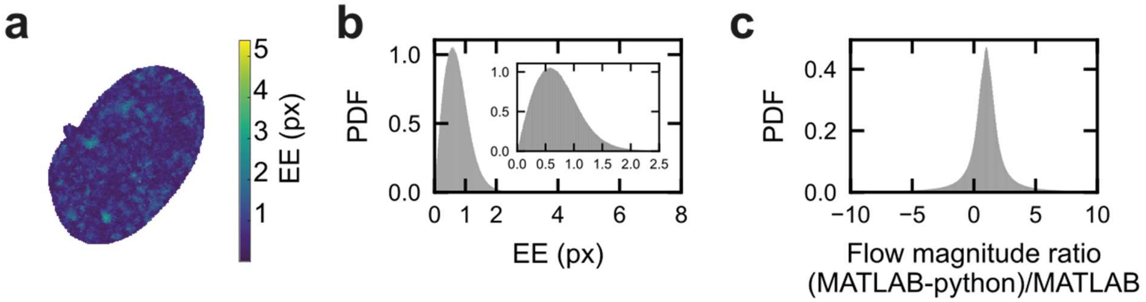


**Figure S1 | Comparison of the Optical Flow implementation in Hi-D (MATLAB) and Hi-D-Py (python).** (**a**) The endpoint error (EE) was spatially mapped in the nucleus for one exemplary time point. (**b**) The endpoint error (EE) between flow fields (the inset shows a magnified region from 0 to 2.5 px) and (**c**) the ratio of the flow field magnitude between MATLAB and python over 149 flow fields. Values larger than 0 indicate pixels where the python implementation underestimates the flow magnitude compared to the MATLAB implementation (n=149 flow fields).

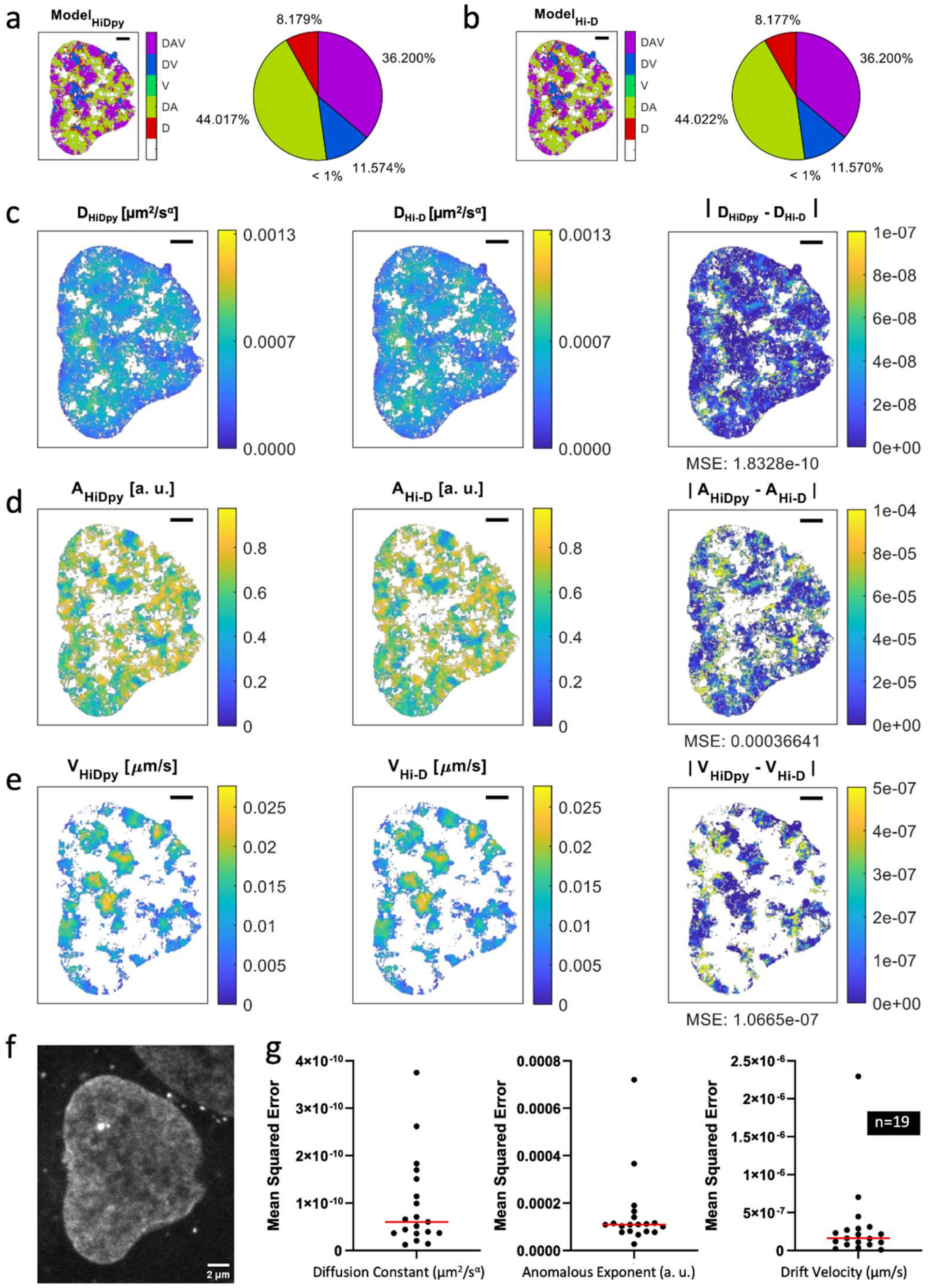


**Figure S2 | Comparison between the Bayesian implementation in Hi-D (MATLAB) and Hi-D-Py (python).** SiR-DNA-stained U2OS cells with Serum were imaged with $\Delta t$ = 200 ms. Trajectories estimated by Hi-D (Matlab) without using denoising were used as input for the Bayesian modules of Hi-D and Hi-D-Py implementations. Bayesian inference of trajectory MSD's results from a representative cell are presented as maps of the selected models (**a**-**b**), diffusion constant (**c**), anomalous exponent (**d**), and drift velocity (**e**). Anomalous exponent fitting was limited between 0 to 1. Differences between coefficients are shown as maps (right **c-e**). Raw representative cell is shown (**f**). Mean squared errors (MSE) are estimated (n = 19 cells).

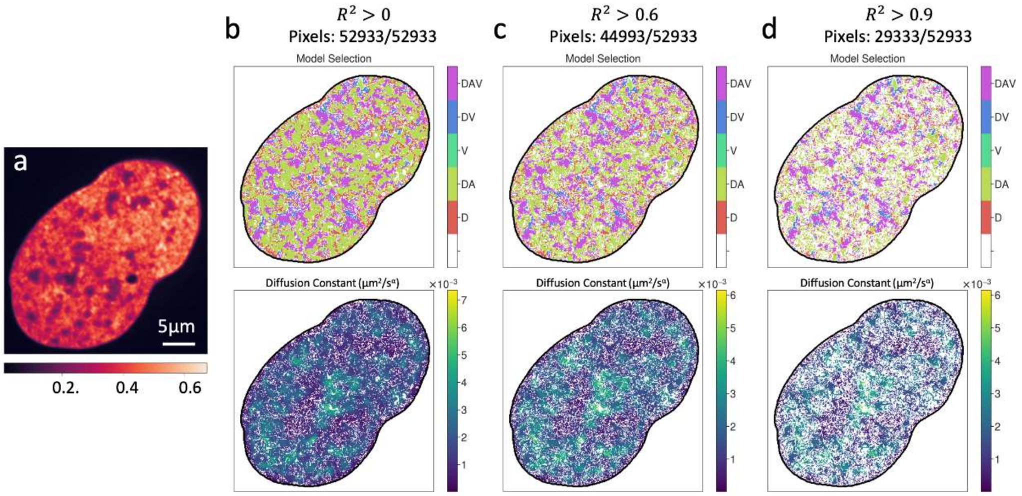


**Figure S3 | Influence of R² of least squares fitting on diffusion constant mapping in Hi-D-Py.** H2B-GFP-stained nuclei of U2OS cells imaged with Δt = 200 ms (a). Model and diffusion coefficient maps were filtered using the estimated R² of the least square fitting part of the Bayesian inference module.

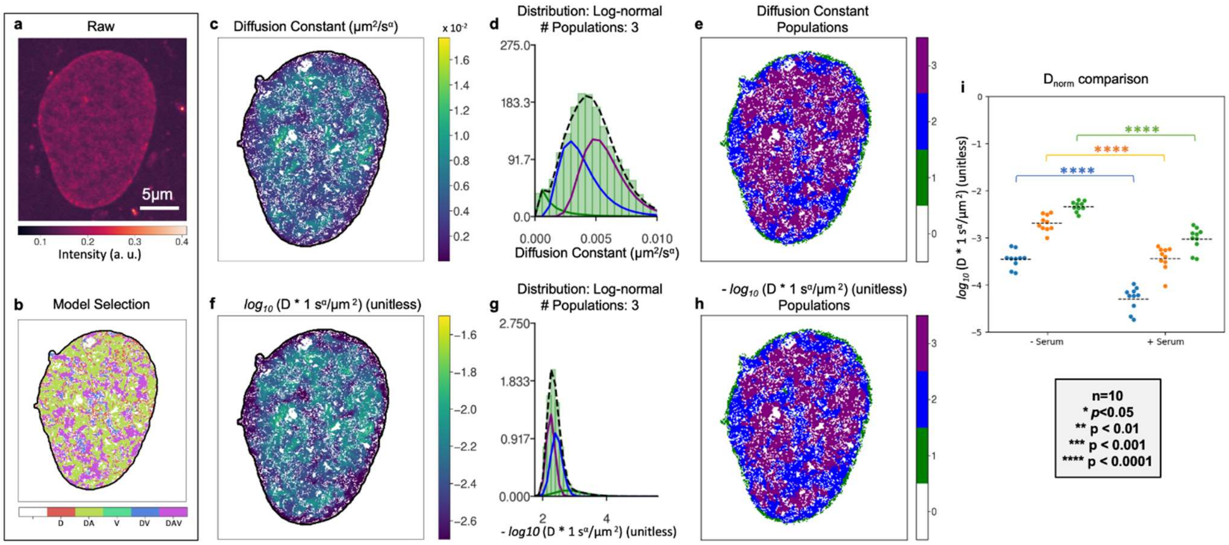


**Figure S4 | Normalization of diffusion constant coefficients.** SiR-DNA-stained U2OS cells without serum were imaged with a time interval (Δt) = 200 ms. Optical flow estimation, Bayesian inference, and diffusion constant normalization were performed using Hi-D-Py. The diffusion-normalized constant ($D_{norm}$) is defined as $D_{norm} = \log_{10}(D \cdot 1\ s^{\alpha}/\mu m^2)$. Representative maps are displayed for a single cell: raw (**a**) and model selection (**b**). Diffusion constant analysis map (**c**) and results from Gaussian Mixture Model (GMM) deconvolution (histograms (**d**) and populations' map (**e**)) are presented. Normalized diffusion constant map (**f**) and results from GMM deconvolution (histograms (**g**) and populations' map (**h**)) are shown. (**i**) Comparison of the normalized diffusion constant averages of SiR-DNA-stained U2OS cells is shown between serum-starved and serum-stimulated conditions (n=10 cells per condition). Average values per population and cell are depicted. Statistically, t-test was performed using the python library scipy.

### References


1. Saxton, M. J. & Jacobson, K. Single-particle tracking: Applications to Membrane Dynamics. *Annual Review of Biophysics and Biomolecular Structure* **26,** 373–399 (1997).
2. Seber, G. & Wild, C. *Nonlinear regression* (John Wiley & Sons, 2003).
3. Wit, E., Heuvel, E. v. d. & Romeijn, J.-W. 'All models are wrong...': an introduction to model uncertainty. *Statistica Neerlandica* **66,** 217–236 (2012).
4. Sun, D., Roth, S. & Black, M. J. A quantitative analysis of current practices in optical flow estimation and the principles behind them. *International Journal of Computer Vision* **106,** 115–137 (2014).
5. Shaban, H. A., Barth, R. & Bystricky, K. Formation of correlated chromatin domains at nanoscale dynamic resolution during transcription. *Nucleic acids research* **46,** e77–e77 (2018).
